# Estradiol-inducible AvrRps4 expression reveals distinct properties of TIR-NLR-mediated effector-triggered immunity

**DOI:** 10.1101/701359

**Authors:** Bruno Pok Man Ngou, Hee-Kyung Ahn, Pingtao Ding, Amey Redkar, Hannah Brown, Yan Ma, Mark Youles, Laurence Tomlinson, Jonathan DG Jones

## Abstract

Plant nucleotide-binding domain, leucine-rich repeat receptor (NLR) proteins play important roles in recognition of pathogen-derived effectors. However, the mechanism by which plant NLRs activate immunity is still largely unknown. The paired Arabidopsis NLRs RRS1-R and RPS4, that confer recognition of bacterial effectors AvrRps4 and PopP2, are well studied, but how the RRS1/RPS4 complex activates early immediate downstream responses upon effector detection is still poorly understood. To study RRS1/RPS4 responses without the influence of cell-surface receptor immune pathways, we generated an Arabidopsis line with inducible expression of effector AvrRps4. Induction does not lead to hypersensitive cell death response (HR) but can induce electrolyte leakage, which often correlates with plant cell death. Activation of RRS1 and RPS4 without pathogens cannot activate mitogen-associated protein kinase cascades, but still activates upregulation of defense genes, and therefore resistance against bacteria.

**Highlight:** Inducible expression of AvrRps4 activates RRS1/RPS4-mediated effector-triggered immunity without the presence of pathogens, allowing us to characterise downstream immune responses triggered by TIR-NLRs without cell-surface receptor-mediated immunity.

## Introduction

To investigate plant immunity, researchers routinely conduct pathogen inoculations on plants in a controlled environment. Upon pathogen attack, plants activate innate immune responses via both membrane-associated and intracellular receptors, which makes it difficult to unravel the distinct contribution of each component. Most plasma-membrane localized receptors perceive conserved pathogen-associated molecular patterns (PAMPs) or host-cell-derived damage-associated molecular patterns (DAMPs) and activate PAMP-triggered immunity (PTI) or DAMP-triggered immunity (DTI). Plant intracellular immune receptors belong to a family of nucleotide-binding leucine-rich repeat (NB-LRR) proteins, also known as NLRs. NLRs recognize pathogen effectors and activate effector-triggered immunity (ETI), which often leads to accumulation of reactive oxygen species (ROS) and a hypersensitive cell death response (HR). Most plant NLRs carry either coiled-coil (CC) or Toll/interleukin-1 receptor (TIR) N-terminal domains. Both CC and TIR domains are believed to function in signaling upon activation of NLRs, but the detailed mechanisms are unknown. Many CC-NLRs localize at and function in association with the plasma membrane, whereas TIR-NLRs can function in diverse locations, including the nucleus. Regardless of the distinct localization patterns between CC- and TIR-NLRs, their downstream outputs culminate in elevated resistance, but have never been directly compared side-by-side. To study the specific immune outputs generated by ETI, inducible expression tools have been applied (McNellis *et al.*, 1998; Tornero *et al.*, 2002; Allen *et al.*, 2004; Porter *et al.*, 2012).

In Arabidopsis, functionally paired NLRs RRS1-R and RPS4 confer resistance against a soil-borne bacterial pathogen *Ralstonia solanacearum* through the recognition of an effector PopP2 secreted via Type III secretion system and a hemibiotrophic ascomycetous fungal pathogen *Colletotrichum higginsianum* (Narusaka *et al.*, 2009). They can also confer resistance against bacteria *Pseudomonas syringae* pv. *tomato* DC3000 carrying AvrRps4, an effector protein from *Pseudomonas syringae* pv. *pisi*, causing bacterial blight in *Pisum sativum* (pea) (Sohn *et al.*, 2009; Narusaka *et al.*, 2009). Previously, it was known that the 135th to 138th residues of AvrRps4, KRVY, are required for the recognition of AvrRps4 by RRS1 and RPS4 (Sohn *et al.*, 2009). Crystal structure information of RRS1 and RPS4 on the TIR domains and the co-crystal structures between RRS1-R WRKY domain and effector PopP2 have indicated some structural basis of how RRS1/RPS4 have been activated (Williams *et al.*, 2014; Zhang *et al.*, 2017). However, it is still unknown how the protein complex assembles and functions.

Here we report tools for studying the immune complex of RRS1-R and RPS4 *in vivo*. We established a set of transgenic Arabidopsis lines to study RRS1/RPS4-mediated ETI in the absence of pathogens. Using these lines, we show that some but not all immune outputs induced by the conditionally expressed AvrRps4 resemble other reported effector-inducible lines.

## Materials and Methods

### Plant material and growth conditions

*Arabidopsis thaliana* accessions Wassilewskija-2 (Ws-2) and Columbia-0 (Col-0) were used as wild type in this study. The *eds1-2* mutant used has been described previously (Falk *et al.*, 1999). Seeds were sown on compost and plants were grown at 21°C with 10 hours under light and 14 hours in dark, and at 70% humidity. Tabaco plants were grown at 22°C with 16 hours under light and 8 hours in dark, and at 80% constant humidity. The light level is approximately 180-200 μmols with fluorescent tubes.

### FastRed selection for transgenic Arabidopsis

Seeds harvested from the Agrobacteria-transformed Arabidopsis are resuspended in 0.1% Agarose and exposed under fluorescence microscope with DsRed (red fluorescent protein) filter. Seeds with bright red fluorescence are selected as the positive transformants.

### GUS staining

*Nicotiana benthamiana* (*N. b.*) leaves were infiltrated with Agrobacteria carrying constructs with β-glucuronidase (GUS) reporter gene expressed under selected Arabidopsis promoters (Table S1). Leaves were collected at 2 days post infiltration (dpi), and vacuum-infiltrated with GUS staining buffer (0.1 M sodium phosphate pH 7.0, 10 mM EDTA pH 7.0, 0.5 mM K_3_Fe(CN)_6_, 0.5 mM K_4_Fe(CN)_6_, 0.76 mM 5-Bromo-4-chloro-3-indolyl-β-D-glucuronide cyclohexylamine salt or X-Gluc, 0.04% Triton X-100). After vacuum-infiltration, the leaves were incubated at 37°C overnight in the dark. The leaves were rinsed with 70% ethanol until the whole leaf de-stains to a clear white.

### Immunoblotting

*N. b.* leaves were infiltrated with Agrobacteria carrying our stacking constructs (Table S2). At 2 dpi, same leaves were infiltrated with either DMSO or 50 μM β-estradiol (E2) diluted in water. Samples were collected at 6 hpi of DMSO or E2 treatment, and snap-frozen in liquid nitrogen. Proteins were extracted using GTEN buffer (10% glycerol, 25 mM Tris pH 7.5, 1 mM EDTA, 150 mM NaCl) with 10 mM DTT, 1% NP-40 and protease inhibitor cocktail (cOmplete™, EDTA-free; Merck). For *Arabidopsis* seedlings, seedlings grown for 8 days after germination were treated with DMSO or E2 with indicated time points and snap-frozen in liquid nitrogen. After centrifugation at 13,000 rpm for 15 minutes to remove cell debris, protein concentration of each sample was measured using the Bradford assay (Protein Assay Dye Reagent Concentrate; Bio-Rad). After normalization, extracts were incubated with 3× SDS sample buffer at 95°C for 5 minutes. 6% SDS-PAGE gels were used to run the protein samples. After transferring proteins from gels to PVDF membranes (Merck-Millipore) using Trans-Blot Turbo System (Bio-Rad), membranes were immunoblotted with HRP-conjugated Flag antibodies (Monoclonal ANTI-FLAG^®^ M2-Peroxidase HRP antibody produced in mouse, A5892; Merck-Millipore), HRP-conjugated HA antibodies (12013819001; Merck-Roche) or Phospho-p44/42 MAPK (Erk1/2) (Thr202/Tyr204) (D13.14.4E) XP® Rabbit monoclonal antibody (4370; Cell Signaling Technology). Anti-Rabbit IgG (whole molecule)–Peroxidase antibody produced in goat (A0545; Merck-Sigma-Aldrich) was used as secondary antibody following the use of Phospho-p44/42 MAPK antibody.

### Bacterial growth assay

*Pseudomonas syringae* pv. *tomato* strain DC3000 carrying pVSP61 empty vector was grown on selective King’s B (KB) medium plates containing 15% (w/v) Agar, 25 μg/ml rifampicin and 50 μg/ml kanamycin for 48 h at 28°C. Bacteria were harvested from the plates, resuspended in infiltration buffer (10 mM MgCl_2_) and the concentration was adjusted to an optical density of 0.001 at 600 nm (OD_600_=0.001, representing approximately 5×10^5^ colony forming units [CFU] ml^−1^). Bacteria were infiltrated into abaxial surfaces of 5-week-old Arabidopsis leaves with a 1-ml needleless syringe. For quantification, leaf samples were harvested with a 6-mm-diameter cork borer (Z165220; Merck-Sigma-Aldrich), resulting in leaf discs with an area of 0.283 cm². Two leaf discs per leaf were harvested as a single sample. For each condition, four samples were collected immediately after infiltration as ‘day 0’ samples to ensure no significant difference introduced by unequal infiltrations and six samples were collected at 3 dpi as ‘day 3’ samples to compare the bacteria growth between different genotypes, conditions and treatments. For ‘day 0’, samples were ground in 200 μl of infiltration buffer and spotted (10 μl per spot) on selective KB medium agar plates to grow for 48 h at 28°C. For ‘day 3’, samples were ground in 200 μl of infiltration buffer, serially diluted (5, 50, 500, 5000 and 50000 times) and spotted (6 μl per spot) on selective KB medium agar plates to grow for 48 h at 28°C. The number of colonies (CFU per drop) was monitored and bacterial growth was represented as in CFU cm^−2^ of leaf tissue. All results are plotted using ggplot2 in R (Wickham, 2009), and detailed statistics summary can be found in the supplemental materials.

### HR assay in Arabidopsis

*Pseudomonas fluorescens* engineered with a type III secretion system (Pf0-1 ‘EtHAn’ strains) expressing one of wild-type or mutant effectors, AvrRps4, AvrRps4^KRVY135-138AAAA^, PopP2, PopP2^C321A^, AvrRpt2 or pVSP61 empty vector were grown on selective KB plates for 24 h at 28°C (Thomas *et al.*, 2009; Sohn *et al.*, 2014). Bacteria were harvested from the plates, resuspended in infiltration buffer (10 mM MgCl_2_) and the concentration was adjusted to OD_600_= 0.2 (10^8^ CFU ml^−1^). The abaxial surfaces of 5-week-old Arabidopsis leaves were hand infiltrated with a 1-ml needleless syringe. Cell death was monitored 24 h after infiltration.

### Electrolyte leakage assay

Either 50 μM E2 or DMSO were hand infiltrated in 5-week-old Arabidopsis leaves with a 1-ml needleless syringe for electrolyte leakage assay. Leaf discs were taken with a 2.4-mm-diameter cork borer from infiltrated leaves. Discs were dried and washed in deionized water for 1 hour before being floated on deionized water (15 discs per sample, three samples per biological replicate). Electrolyte leakage was measured as water conductivity with a Pocket Water Quality Meters (LAQUAtwin-EC-33; Horiba) at the indicated time points. All results are plotted using ggplot2 in R (Wickham, 2009), and detailed statistics summary can be found in the supplemental materials.

### Trypan blue staining

Either 50 μM E2 or DMSO were hand infiltrated in 5-week-old Arabidopsis leaves with a 1-ml needleless syringe for trypan blue staining. 6 leaves per sample were collected 24 hours after infiltration. Leaves were boiled in trypan blue solution (1.25 mg/ml trypan blue dissolved in 12.5% glycerol, 12.5% phenol, 12.5% lactic acid and 50% ethanol) in a boiling water bath for 1 min and de-stained by chloral hydrate solution (2.5 g/ml). De-stained leaves were mounted, and pictures were taken under on Leica fluorescent stereomicroscope M165FC. All images were taken with identical settings at 2.5x magnification. Scale bar=0.5mm.

### Gene expression measurement by reverse transcription-quantitative polymerase chain reaction (RT-qPCR)

For gene expression analysis, RNA was isolated from 5-week-old Arabidopsis leaves and used for subsequent RT-qPCR analysis. RNA was extracted with Quick-RNA Plant Kit (R2024; Zymo Research) and treated with RNase-free DNase (4716728001; Merck-Roche). Reverse transcription was carried out using SuperScript IV Reverse Transcriptase (18090050; ThermoFisher Scientific). qPCR was performed using a CFX96 Touch^TM^ Real-Time PCR Detection System. Primers for qPCR analysis of *Isochorismate Synthase1* (*ICS1*), *Pathogenesis-Related1* (*PR1*), *AvrRps4* and *Elongation Factor 1 Alpha* (*EF1*α) are listed in Table S4. Data were analyzed using the double delta Ct method (Livak and Schmittgen, 2001). All results are plotted using ggplot2 in R (Wickham, 2009), and detailed statistics summary can be found in the supplemental materials.

### Confocal laser scanning microscopy (CLSM) imaging

Transgenic plant materials were imaged with the Leica DM6000/TCS SP5 confocal microscopy (Leica Microsystems) for confirmation of expression of inducible AvrRps4 fused with monomeric yellow-green fluorescent protein, mNeonGreen or mNeon (Shaner *et al.*, 2013). Roots from 3-week-old Arabidopsis seedlings were sprayed with 50 μM E2 and imaged at 1 day post spray. Fluorescence of mNeon was excited at 500 nm and detected at between 520 and 540 nm. CLSM images of root cells from Arabidopsis seedlings are recorded via the camera. The images were analyzed with the Leica application Suite and Fiji software (Schindelin *et al.*, 2012).

### Co-immunoprecipitation

Arabidopsis transgenic seedlings, and the background ecotype Col-0 grown for 7 days after germination (DAG) were treated with 0.1% DMSO or 50μM E2 for 3 hours. Proteins from seedlings were extracted using IP buffer (10% Glycerol, 50mM Tris-Cl pH 6.8, 50mM KCl, 1mM EDTA, 5Mm MgCl_2_, 1% NP-40, 10mM DTT, 1mM dATP). Crude extract of the seedlings was centrifuged and supernatants were incubated with Anti-HA-conjugated beads (EZviewTM Red Anti-HA Affinity Gel; E6779; Sigma). A small portion of supernatants were taken for input samples. At 2 hours after incubation of the extract with beads, beads were washed three times with IP buffer containing 0.1% NP-40. Proteins bound to beads were eluted by boiling the beads with SDS sample buffer. Immunoblotting of the input and eluted samples were performed as described above.

## Results

### RRS1 over-expression can compromise RPS1/RPS4 function

Overexpression of *RPS4* leads to autoimmunity and dwarfism under standard growth condition (see methods) (Heidrich *et al.*, 2013). This autoimmunity is both temperature- and RRS1-dependent. In contrast, elevated expression of *RRS1-R* from ecotype Ws-2 in Col-0, an ecotype expressing a dominant allele of *RRS1-S*, does not trigger auto-immunity (Huh et al 2017). Furthermore, high level RRS1-R expression does not confer recognition of effector PopP2 (Fig 1). Overexpression of *RRS1* in an *RPS4* overexpression line attenuates dwarfism and autoimmunity (Huh *et al.*, 2017). Here, we found that only simultaneously over-expressing both *RRS1* and *RPS4* can lead to the gain-of-recognition of PopP2 in the susceptible ecotype Col-0 (Fig 1). Thus, we propose that a balanced protein expression of RRS1 and RPS4 is required for both suppressing autoimmunity and functional recognition of the corresponding effectors.

**Fig. 1.**
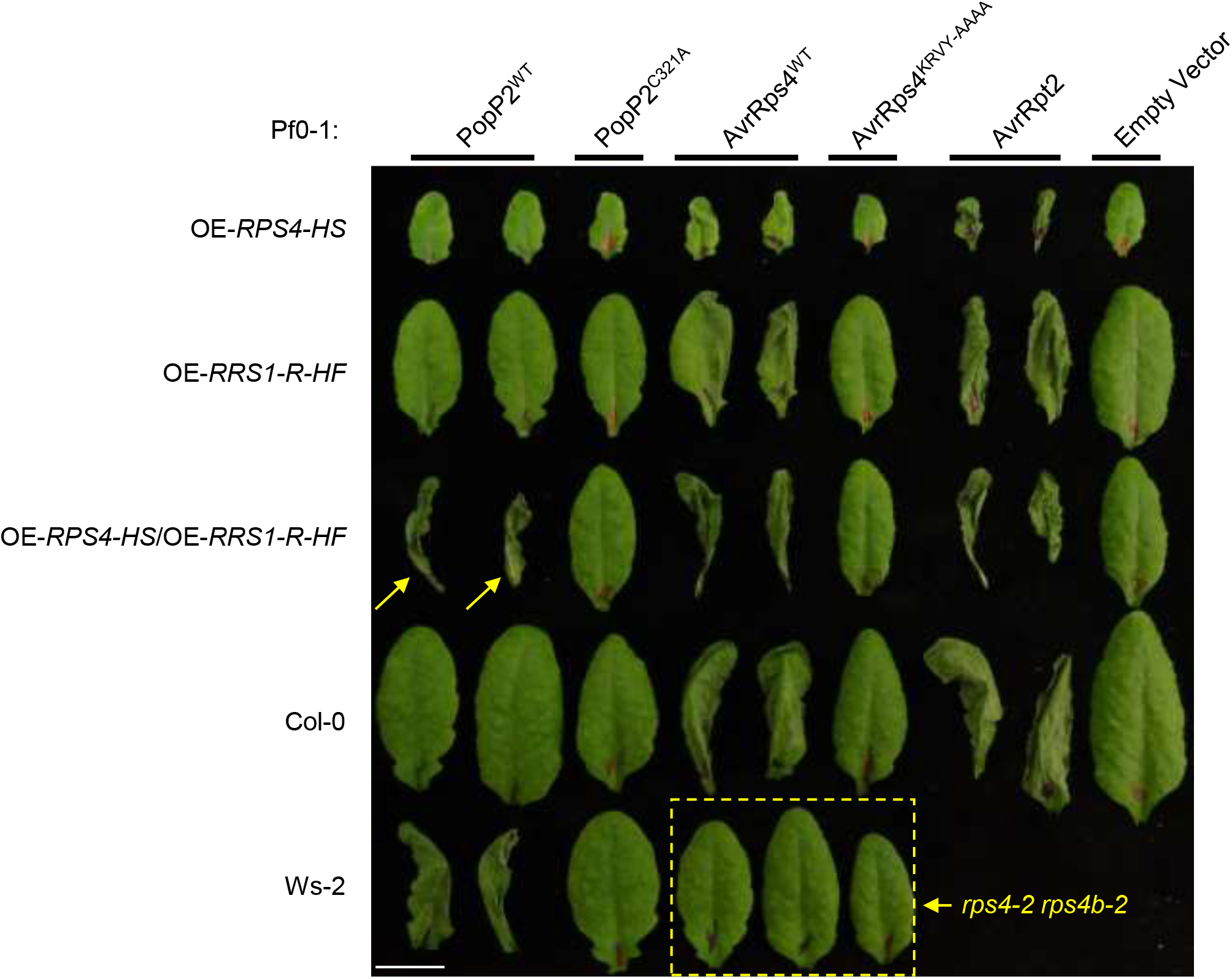
Over-expression of RPS4 and RRS1-R reconstruct the recognition of PopP2 in Col-0. Arabidopsis transgenic lines overexpressing RPS4 (OE-RPS4-HS), RRS1-R (OE-RRS1-R-HF) or both generated by crossing (OE-RPS4-HS/OE-RRS1-R-HF) in the Col-0 background, with Col-0 and Ws-2 accession were tested for hypersensitive response (HR). 5-week old leaves were infiltrated with Pseudomonas fluorescence (Pf) 0-1 strains carrying empty vector (EV), wild-type (WT) AvrRps4, mutant AvrRps4^KRVY-AAAA^, WT PopP2, mutant PopP2^C321A^, and WT AvrRpt2. Leaves were collected 1 day post infiltration (dpi) for imaging. Scale bar = 1cm. Yellow arrows indicate reconstructed PopP2 recognition of Col-0 background overexpressing RRS1-R and RPS4. Yellow dashed box highlights loss of AvrRps4 recognition in the double mutant rps4-2 rps4b-2. Infiltration of EV and AvrRpt2 serve as negative and positive controls of HR, respectively.

### A survey of leaf-expressed genes reveals promoters for moderate and balanced expression levels of RRS1 and RPS4

Genome-wide expression profiling has revealed numerous genes altered by PTI alone or PTI plus ETI at early time points of RRS1/RPS4-mediated immune activation (Sohn *et al.*, 2014). This analysis also enabled the discovery of genes that are moderately and constitutively expressed without changing their transcript abundance during immune activation. In plants, gene expression patterns and levels are usually specified by their promoters. Based on the endogenous expression relative transcript abundance in the ‘stable gene set’, we selected six promoters with ‘moderate’ expression (Table S1). We define the ‘moderate’ expression based on two criteria: (1) the gene transcript abundance with those promoters are at least 100 times more than the endogenous transcript abundance of *RRS1* and *RPS4*; (2) the gene transcript abundance with those promoters is lower than that with the 35S promoter. The selected genes encode proteins that are involved in essential biological processes that we expect to be expressed in most mesophyll cells, including a delta-tonoplast intrinsic protein (our name; At1, locus identifier AT3G16240, protein symbol name TIP2-1), a ribosomal protein S16 (At2, AT4G34620, RPS16-1), a cysteine synthase isomer CysC1 (At3, AT3G61440, CYSC1), a photosystem II subunit Q (At4, AT4G21280, PSBQ1), a xyloglucan endotransglucosylase/hydrolase 6 (At5, AT5G65730, XTH6), and a ubiquitin-like protein 5 (At6, AT5G42300, UBL5) (Table S1).

To test the strength of the selected Arabidopsis promoters (pAt1-pAt6) for driving gene expression *in planta*, constructs were designed and generated to use them to express β-glucuronidase (GUS) (pAt:GUS). Agrobacterium strains carrying each pAt:GUS construct was infiltrated in tobacco leaves with the infiltration buffer as negative control and GUS expressed under the CaMV 35S promoter (35S:GUS) as positive control. GUS expressed under pAt4 shows similar level of activity to that with 35S, whereas GUS activities detected from other pAt promoters are significantly weaker (Fig S1).

### A T-DNA construct expresses RPS4, RRS1 and inducible AvrRps4

We designed a binary vector to reconstruct the effector ligand AvrRps4 and its receptors RRS1 and RPS4, using the Golden Gate Modular Cloning Toolbox (Fig 2A) (Engler *et al.*, 2014). We chose moderate and balanced promoters pAt2 and pAt3 from our promoter survey experiment for expressing RRS1 and RPS4, respectively. We have also cloned *RRS1-R* full-length coding sequences (CDS) from Ws-2 and *RPS4* full-length CDS from Col-0 for the expression of RRS1-R and RPS4 proteins. We chose synthetic C-terminal-fusion epitope tags His_6_-TEV-FLAG_3_ (HF) and HA_6_ for detecting RRS1 and RPS4 protein expressions, respectively (Fig 2A, Table S2) (Gauss *et al.*, 2005; Soleimani *et al.*, 2013). We have used an E2-inducible system for AvrRps4 expression (Zuo *et al.*, 2000). We named this multi-gene stacking binary construct ‘Super ETI’, or SETI. All restriction enzyme sites for BsaI and BpiI in modules for promoters, CDSs for genes or epitope tags and the terminators were synonymously eliminated (Fig 2A, Table S1). More detailed information for the cloning can be found in supplemental materials.

**Fig 2.**
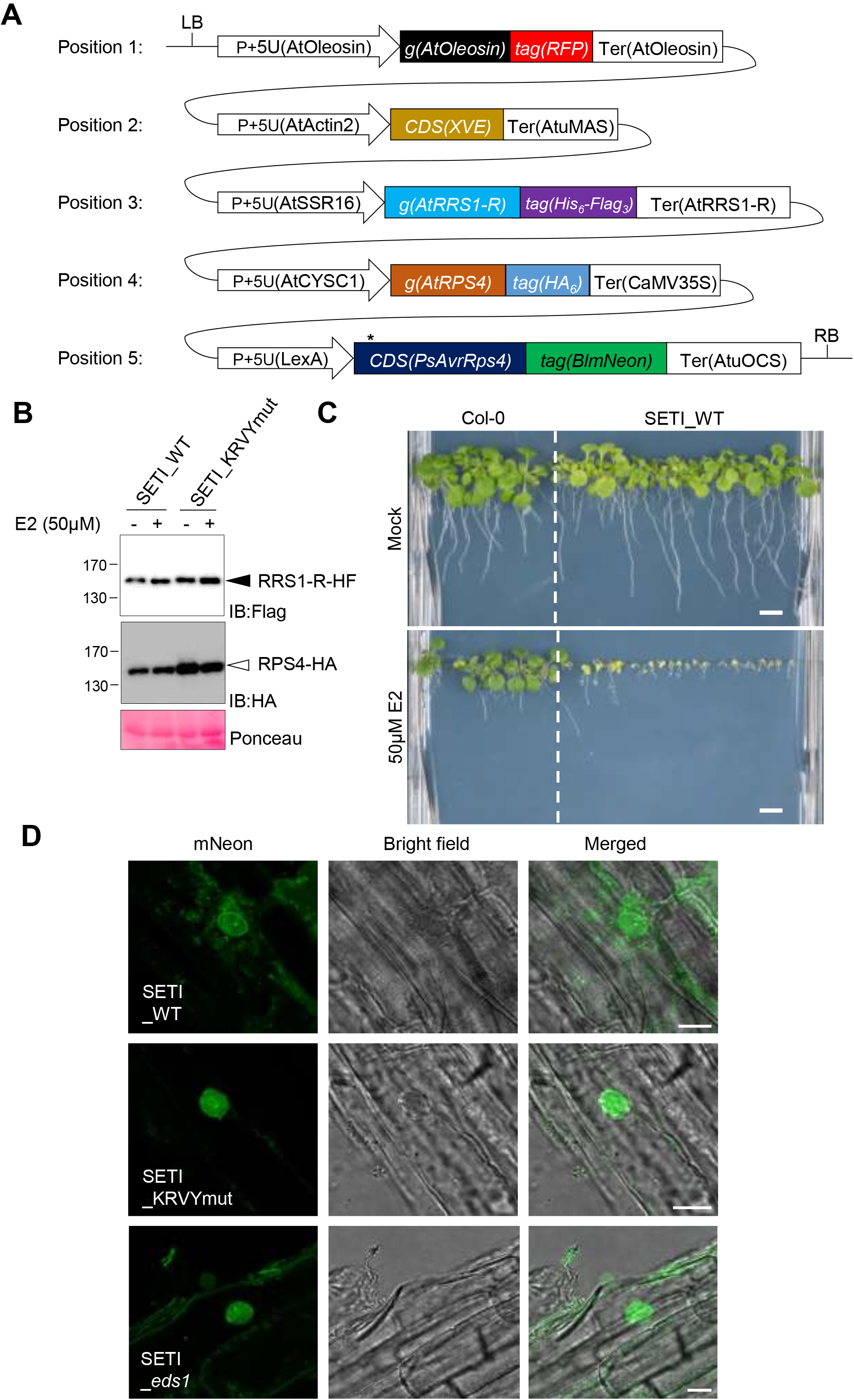
Single T-DNA expresses RRS1-R-HF, RPS4-HA and inducible wild-type AvrRps4 or AvrRps4 mutant variants (A) Illustrative layout of the SUPER-ETI (SETI) construct. There are five individual expression units or Golden Gate Level 2 positional components listed, which are indicated position 1 to position 5. Position 1; expression unit of the FastRed selection marker (Shimada et al. 2010). Position 2, 5; chimeric transactivator XVE (LexA-VP16-ER) and the corresponding LexA inducible system to express AvrRps4 or its mutant variants under the control of β-estradiol (E2) treatment. Position 3, 4; full-length RRS1-R and RPS4 proteins with epitope tags His_6_-Flag_3_ and HA_6_, respectively. All cloning details can be found in Methods and Materials. All individual units used for construct assembly can be found in the Supplemental Table 2 and 3. (B) Protein accumulation of RRS1-R-HF (IB:Flag, black arrowhead) and RPS4-HA (IB:HA, white arrowhead) of SETI lines expressing AvrRps4 (SETI_WT) or mutant AvrRps4 KRVY-AAAA (SETI_KRVYmut). Seedlings were grown in liquid culture and induced with 50μM E2 for 2 hours at 7 days after germination (DAG). Ponceau staining of Rubisco large subunits were used as loading control. (C) Seedling phenotype of SETI Arabidopsis transgenic line at 14 DAG in GM media containing Mock (0.1% DMSO) or 50μM E2. Col-0 was sown as control for the effect of E2 on seedling growth. Scale bar = 0.5cm (D) Confocal images of SETI_WT, SETI_KRVYmut, SETI_*eds1* root cells expressing AvrRps4-mNeon and AvrRps4^KRVY-AAAA^–mNeon induced by 50μM E2 for 24h. mNeon channel shows nucleo-cytoplasmic localization of AvrRps4-mNeon and AvrRps4^KRVY-AAAA^–mNeon. Bright field channel and merged image of mNeon and Bright field channel are shown together. Bars = 10 μm.

To verify the SETI construct, we used a transient expression system in *Nicotiana benthamiana* by infiltrating Agrobacteria that deliver the SETI T-DNA. Protein accumulation of RRS1-R-HF and RPS4-HA was detected (Fig S2).

### The single-locus lines carrying the SETI T-DNA show inducible growth arrest

We generated transgenic Arabidopsis lines using the SETI construct expressing wild-type AvrRps4 (SETI_WT). With the FastRed selection module, we have selected several positive SETI_WT lines (see Materials and Methods, Table S2, Fig S3). We confirmed protein expression of RRS1-R-HF and RPS4-HA (Fig 2B). We also tested the expression of inducible AvrRps4-mNeon under fluorescence microscope upon the treatment with E2. mNeonGreen signal was detected at 24 hours post spray on transgenic seedlings, consistent with the mRNA accumulation of *AvrRps4* at 4 hours post E2-infiltration in leaves (Fig 2D, Fig S2C). On E2-containing growth medium, SETI_WT transgenic lines display severe growth arrest (Fig 2C, Fig S3). We selected one of the lines (SETI_WT) for subsequent experiments (Fig 2C, Fig S3).

### RRS1-R and RPS4 form pre-activation complexes in Arabidopsis

The SETI lines enable detection of epitope-tagged RRS1-R and RPS4 (Fig 2B). We investigated in vivo interaction of tagged RRS1-R and RPS4 by co-immunoprecipitation (co-IP). When RPS4-HA was immunoprecipitated using HA beads, we found RRS1-R and RPS4 stay in association with each other both before and 3 hours after the induction of *AvrRps4* expression (Fig 3). There were no significant differences of RRS1-R and RPS4 association upon *AvrRps4* induction. While all previous studies in interactions of RRS1-R and RPS4 was tested only using *N. benthamiana* transient expression system (Huh *et al.*, 2017), generation of SETI line enabled the detection of RRS1-R and RPS4 interaction in its native system in Arabidopsis.

**Fig 3.**
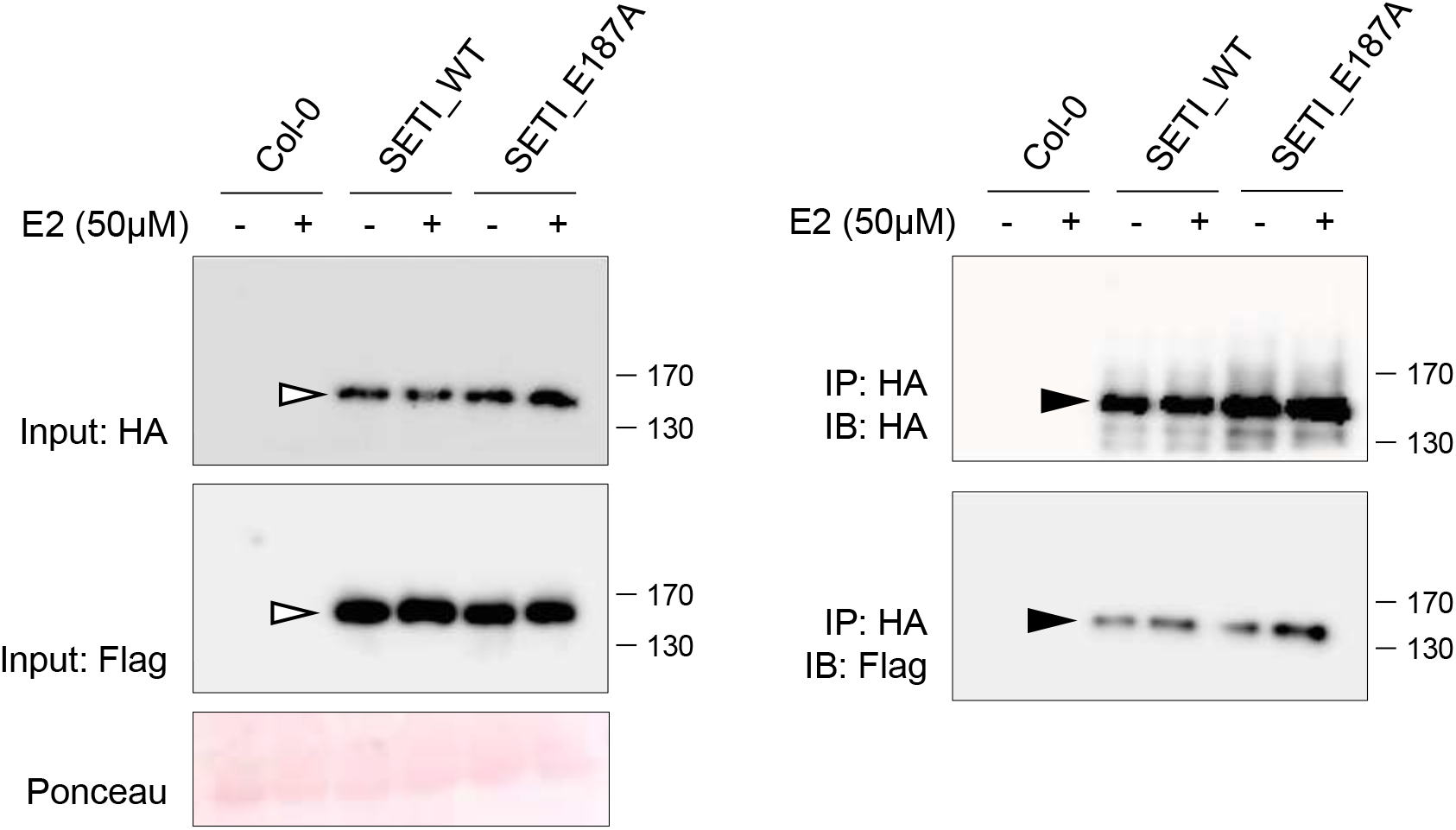
RRS1-R and RPS4 interact *in vivo*. Co-immunoprecipitation of RRS1-R-HF with RPS4-HA. Col-0, SETI_WT, and SETI_E187A seedlings at DAG7 were treated with 50μM E2 for 3hours. Crude extracts were centrifuged and RPS4-HA proteins were immunoprecipitated with Anti-HA-conjugated beads. Immunoprecipitation of RPS4-HA, and co-immunoprecipitation of RRS1-R-HF were determined by immunoblot analysis with HA (IB:HA) or Flag (IB:Flag). Ponceau staining indicates equal loading of the input samples. RRS1-R-HF (black arrowhead), and RPS4-HA (white arrowhead) are indicated.

### Some but not all defense responses are induced by E2 in SETI lines

The induced expression of multiple effectors, such as AvrRpt2, AvrRpm1 and ATR13 can induce cell death or named macroscopic HR in Arabidopsis leaves (McNellis *et al.*, 1998; Tornero *et al.*, 2002; Allen *et al.*, 2004). We therefore tested whether induced expression of AvrRps4 can trigger macroscopic HR in Arabidopsis. As seen in Fig 4A, no HR can be observed after AvrRps4 expression is induced in the SETI leaves. However, only the expression of AvrRps4 but not mutant AvrRps4 (here referred to as AvrRps4_KRVYmut_t_, with residues KRVY 135 to 138 mutated into AAAA) leads to electrolyte leakage (Fig 4C). We also observed slightly stronger trypan blue stains in the SETI leaves treated with E2 compared to mock treatment; suggesting that the expression of AvrRps4 causes microscopic or weak but not macroscopic or strong HR in contrast to other known inducible effector lines (Fig 4B).

**Fig. 4.**
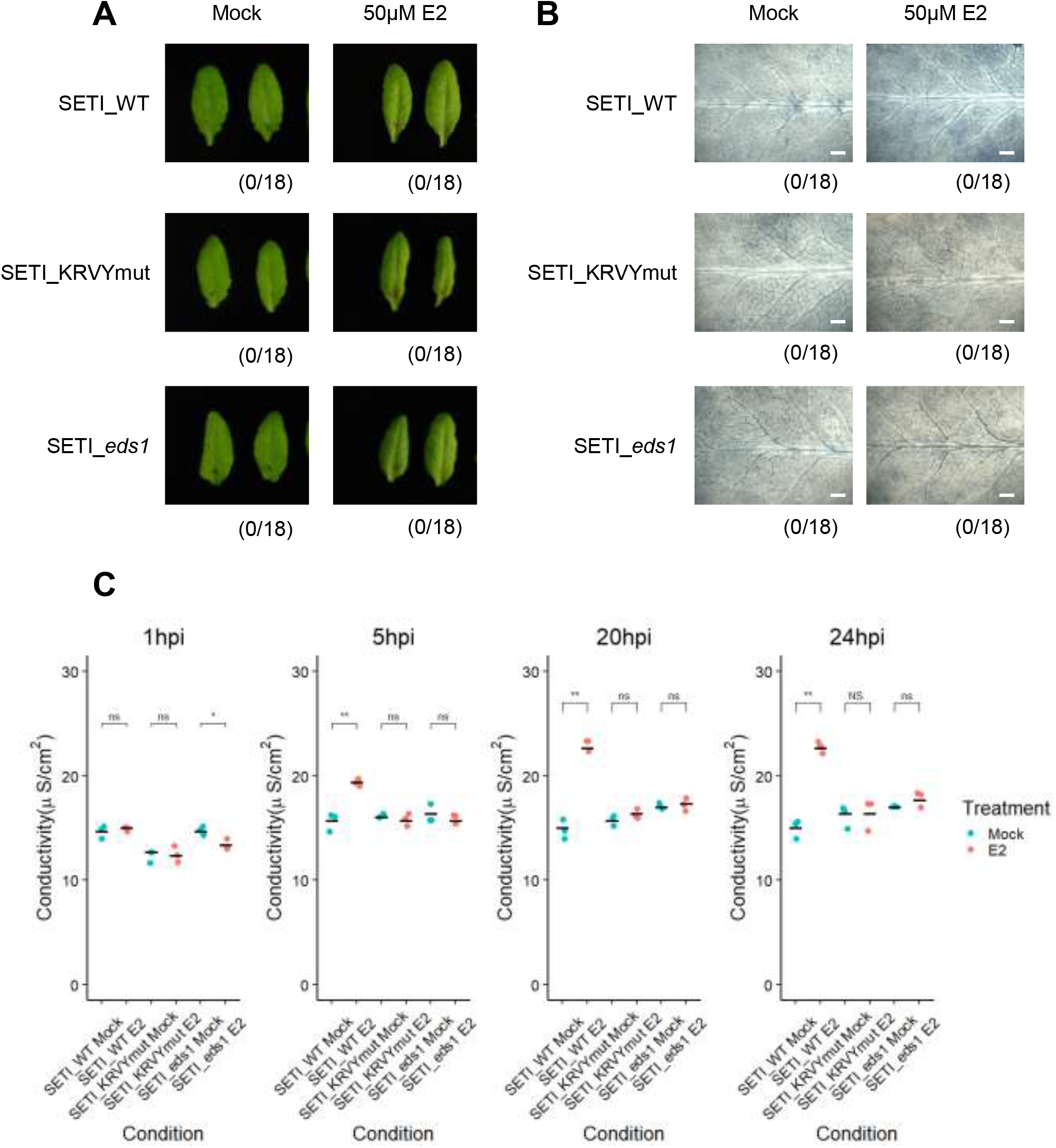
Induced expression of AvrRps4 in *Arabidopsis* cause microscopic but not macroscopic cell death. (A) HR phenotype assay in Arabidopsis. 5-week old SETI_WT, SETI_KRVYmut and SETI_*eds1* leaves were infiltrated with Mock (1% DMSO) or 50μM E2. Images were taken at 1dpi. Numbers indicate the number of leaves displaying cell death from the total number of infiltrated leaves (18 for each genotype and treatment). (B) Trypan blue staining. 5-week old SETI_WT, SETI_KRVYmut and SETI_*eds1* were infiltrated with Mock (1% DMSO) or 50μM E2. Leaves were stained with trypan blue solution at 1dpi. After destaining, leaves were imaged using stereoscopic microscope. Scale bar = 0.5mm (C) Electrolyte leakage assay. 5-week old SETI_WT, SETI_KRVYmut and SETI_*eds1* leaves were infiltrated with Mock (1% DMSO) or 50μM E2. Fifteen leaf discs were collected for each data point. Conductivity was measured at 1, 5, 20 and 24 hours post infiltration (hpi). Each data point represents one technical replicate and three technical replicates are included per treatment and genotype for one biological replicate. Black line represents the mean of the technical replicates. This experiment was repeated three times independently with similar results (Supplemental Figure 2). Significant differences relative to the mock treatment in each genotype was calculated with t-test and the P-values are indicated as ns (non-significant), P > 0.05; *, P < 0.05; **, P < 0.01; ***, P < 0.001.

Salicylic acid induction is another hallmark of ETI (Castel *et al.*, 2019). Enzymes such as Isochorismate Synthase 1 (ICS1), Enhanced Disease Susceptibility 5 (EDS5) and AvrPphB Susceptible 3 (PBS3) are involved in the biosynthesis of salicylic acid and the expression of these genes is also highly induced during ETI (Sohn *et al.*, 2014). The expression of *ICS1* after the *AvrRps4* induction was tested by quantitative real-time PCR. *ICS1* was highly induced 4 hours after the induction of *AvrRps4* by E2 (Fig 5A). In contrast, *Pathogenesis-Related protein 1* (*PR1*) was highly induced only 8 hours after the induction of *AvrRps4* (Fig 5B). This shows that ETI triggered by RRS1/RPS4 is sufficient for the induction of *ICS1* and the biosynthesis of salicylic acid, which subsequently leads to expression of *PR1*.

**Fig. 5.**
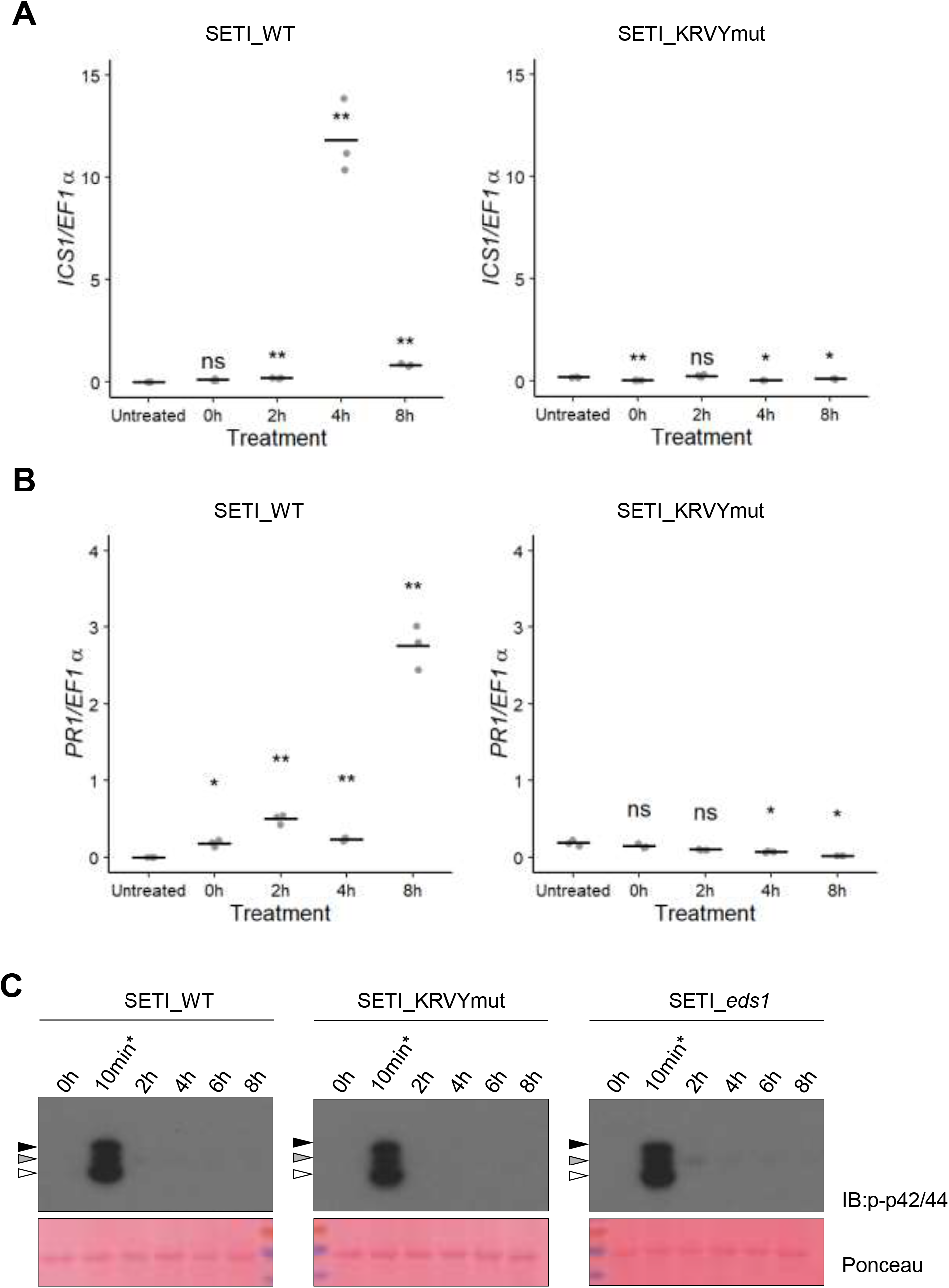
Induced expression of AvrRps4 in *Arabidopsis* leads to *ICS1* and *PR1* expression, but not MAPK activation. (A and B) *ICS1* (A) and *PR1* (B) expression after induction with E2 for 2, 4, and 8h in SETI (left panel) and SETI_KRVYmut (right panel) leaf samples. 5-week old SETI and SETI_KRVYmut leaves were infiltrated with 50 μM E2. Samples were collected at 0, 2, 4 and 8hpi for RNA extraction and subsequent qPCR. Expression level is presented as relative to *EF1α* expression. Each data point represents one technical replicate. Black line represents the mean of the technical replicates. This experiment was repeated three times independently with similar results. Significant differences relative to the Untreated samples was calculated with t-test and the P-values are indicated as ns (non-significant), P > 0.05; *, P < 0.05; **, P < 0.01; ***, P < 0.001. (C) Activation of MAP kinases in SETI_WT, SETI_KRVYmut and SETI_*eds1* seedlings by E2-induction of effector AvrRps4 or mutant AvrRps4^KRVY-AAAA^. Seedlings grown in liquid culture at 7 dag were treated with 50μM E2 for indicated time points (0, 2, 4, 6, 8h) and collected for samples. SETI_WT, SETI_KRVYmut and SETI_*eds1* seedlings treated with 100nM flg22 for 10 minutes (10min*) were used as positive control. Proteins were extracted from these seedlings and phosphorylated MAP kinases were detected using p-p42/44 antibodies. Arrowheads indicate phosphorylated MAP kinases (black; pMPK6, grey; pMPK3, white; pMPK4/11). Ponceau staining were used as loading control.

Activation of mitogen-activated protein kinases (MAPKs) by PTI has been reported under many cases and happens within a few minutes of the activation of PTI. However, the activation of MAPKs by ETI is slower and lasts longer than PTI-induced MAPK activation (Tsuda et al. 2013). We tested whether the induced expression of *AvrRps4* can lead to MAPK activation in SETI_WT and control lines. In contrast to AvrRpt2-inducible transgenic Arabidopsis plants, induced expression of *AvrRps4* does not activate MAPKs (Fig 5C) (Tsuda et al. 2013).

We further tested if the induction of ETI would elevate resistance. We infiltrated the leaves with E2 or mock solution one day before we infiltrated plants with *Pst* DC3000 (Materials and Methods). SETI_WT plants pre-treated with E2 are more resistant to the bacteria than those pre-treated with mock, while there was no significant difference between E2 and mock pre-treatment in Col-0 (Fig 6).

**Fig. 6.**
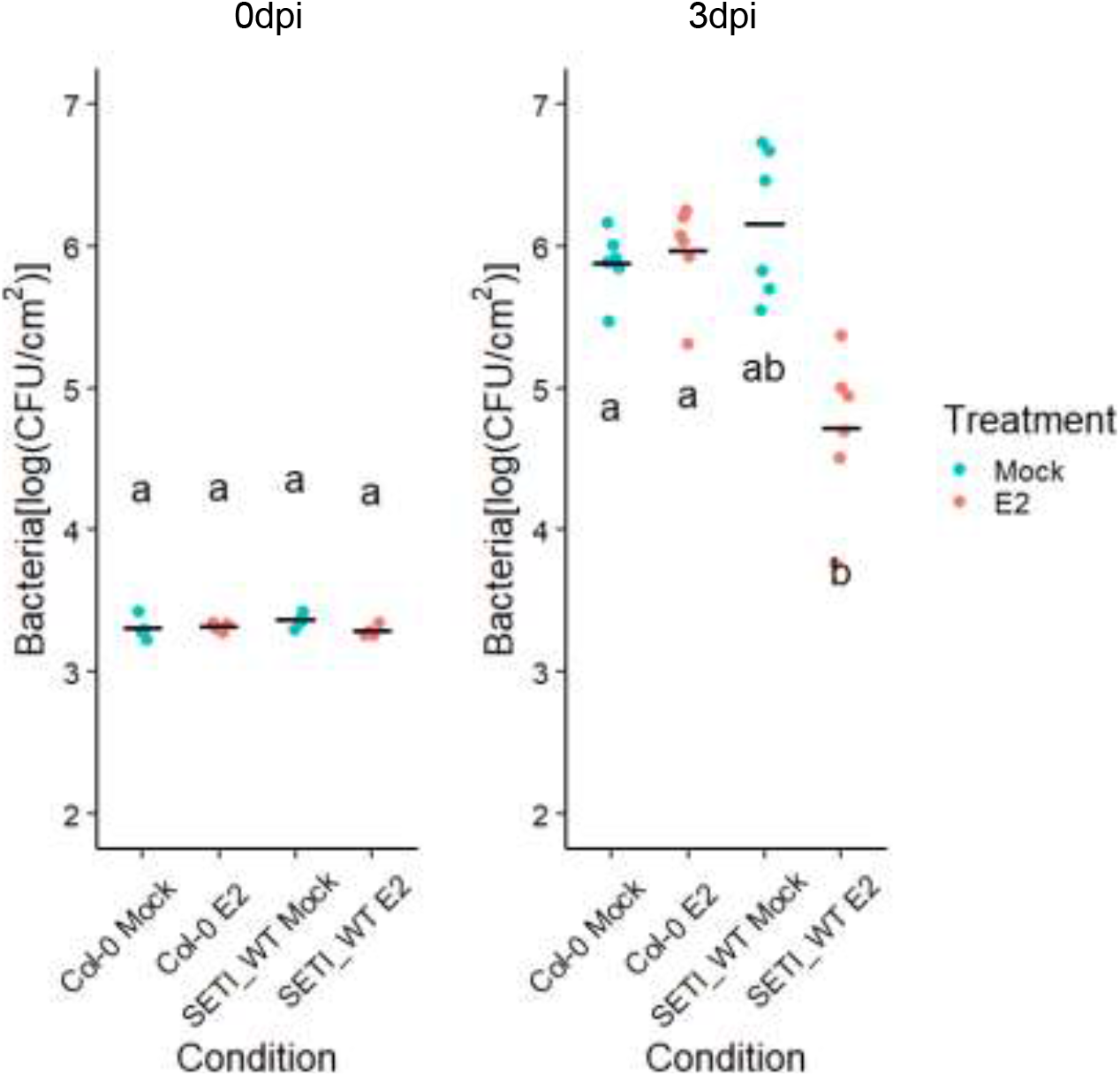
Effector-triggered immunity triggered by the expression of AvrRps4 leads to resistance against *Pseudomonas syringae pv. tomato* strain DC3000. 5-week old SETI_WT and Col-0 leaves were infiltrated with Mock (1% DMSO) or 50μM E2. At 1dpi, leaves were inoculated with *Pst*DC3000 (OD600=0.001). Bacteria in the leaves were then quantified as colony-forming units (CFU) at 0dpi and 3dpi. Each data point represents two leaves collected from one individual plant. Samples from four individual plants were collected for 0 dpi and samples from six individual plants were collected for 3 dpi. Black line represents the mean of the technical replicates. This experiment was repeated three times independently with similar results. Biological significance of the values were determined by one-way ANOVA followed by post hoc TukeyHSD analysis. Letters above the data points indicate significant differences (P<0.05).

## Discussion

To facilitate studying the functional complex of RRS1 and RPS4 *in vivo*, we generated an expression construct of E2-inducible AvrRps4 stacked with epitope-tagged RRS1 and RPS4. To achieve balanced expression levels higher than endogenous expression of *RRS1* and *RPS4*, we surveyed constitutively expressed gene promoters. Here, we report 6 new and tested promoter modules that are compatible with the Golden Gate Modular Cloning toolkit. We used two of the promoters to express *RRS1* and *RPS4*, and we avoided autoimmunity induced by excessive expression of *RPS4* or non-recognition of PopP2 cause by excessive expression of *RRS1-R*. We were also able to generate inducible AvrRps4 expression to activate RRS1/RPS4-mediated ETI under the control of E2 treatment. We thus were able to stack genes for inducible expression of a pathogen effector and its NLR receptors in one construct. In addition, with the epitope tags, we are able to monitor effector-dependent changes in the NLR proteins without interference from using a pathogen effector-delivery system. We could thus express any effectors or pathogen ligands that will trigger immunity in plant cells with the E2-inducible module, and their immune receptors using the same gene stacking strategy.

There are multiple advantages to enabling investigation of ETI without the complication of co-activating PTI. Firstly, we could test the contribution of other genes to ETI activation by introducing mutants into the SETI background, either using conventional crossing or using genome-editing such as CRISPR/Cas9. These lines can also help investigating downstream signaling from plant NLRs. Multiple forward genetic screens have been conducted, but few novel components have been found, and most mutations are either in the NLRs or regulatory elements rather than signaling components (van Wersch *et al.*, 2016). Another plausible explanation is that the signaling path downstream of plants NLRs is very short, but this is debatable, because several significant steps are required for immunity. EDS1, PAD4 and SAG101 are required for TIR-NLR signaling (Falk *et al.*, 1999; Gantner *et al.*, 2019). NRC family proteins in Solanaceae species required for many NLRs, and NRG1/ADR1s in Arabidopsis required for TIR-NLRs and ADR1s for some CC-NLRs (Bonardi *et al.*, 2011; Dong *et al.*, 2016; Wu *et al.*, 2017, 2019; Castel *et al.*, 2019). NRG1s and ADR1s seem to function downstream of EDS1 and may function distinctly with SAG101 and PAD4, respectively (Lapin *et al.*, 2019). SETI lines carry heterologously expressed RRS1-R/RPS4 and also endogenous RRS1-S/RPS4, RRS1B/RPS4B, which together provide three redundant copies of NLR pairs that can recognize AvrRps4. In theory, in an EMS-mutagenesis forward genetic screen to identify suppressors of immunity induced by AvrRps4, there should be a reduced background of mutations in the receptor(s), improving prospects to reveal mutations in genes that are functionally important in NLR signaling and regulation.

With SETI, we are able to assess pure ETI response mediated by the TIR-NLRs, RRS1 and RPS4. E2 induction provoked rapid transcriptional changes in activation of defense genes and also ion leakage. AvrRps4-induced ETI enhanced resistance against bacterial pathogens. However, neither MAPK activation nor macroscopic HR, in contrast to other inducible ETI examples (Tornero *et al.*, 2002; Tsuda *et al.*, 2013). This indicates that outputs of plant NLRs might differ. Both TIR and CC domains alone are sufficient to activate plant immunity. However, whether they signal through similar or different downstream components is still unknown.

In diverse multicellular eukaryotes, immune complexes are assembled into oligomeric complexes to signal downstream. The mammalian inflammasome, assembled in response to bacterial peptide recognition by NAIP proteins and subsequent activation and binding of NLRC4 proteins, is a classic example (Zhang *et al.*, 2015). The plant CC NLR ZAR1 forms an effector-dependent resistosome, which is a pentamer of ZAR1 assembled together with cofactors PBL2 and RKS1 (Wang *et al.*, 2019). The structure of TIR domains implies that activation might require the disassociation of the RRS1 and RPS4 TIR domains and the oligomerization of RPS4 TIR domains (Williams *et al.*, 2014). In SETI lines, RRS1 and RPS4 form a pre-activation complex in the absence of pathogen effector. However, co-IP data cannot distinguish the ratio of which RRS1 and RPS4 bind to each other. It will be interesting to check via various non-denaturing methods if RRS1-R and RPS4 form a dimer or a higher order oligomerization *in vivo,* or whether there is a conformational change in complex upon effector recognition. Furthermore, with the SETI lines generated in this study, we can ask what other co-factors are required for the activation of RRS1-R and RPS4 at native conditions.

The availability of SETI lines also will enable us to study how PTI and ETI interact with each other, especially in the context of RRS1- and RPS4-mediated immunity. Some models have been proposed in discussing on this topic (Tsuda *et al.*, 2009; Cui *et al.*, 2015). From the zig-zag model, PTI and ETI holds in different threshold on activating immunity (Jones and Dangl, 2006). With SETI line, we could specifically ask how physically PTI and ETI can influence each other. A lot of evidence shows that the PTI receptors PRRs usually have very specific post-translational modification events at early time points, there is also some evidence showing ETI can activate somewhat overlapping but different PTMs on immune-related proteins (Withers and Dong, 2017; Kadota *et al.*, 2019). It will be interesting to know how the activation of RRS1/PRS4 leads to the changes of PTMs and how those changes contribute to the robustness of immunity. In addition, transcriptional changes are not the only process reported as the early changes of ETI but also the changes in translations (Meteignier *et al.*, 2017; Xu *et al.*, 2017). Both work using inducible AvrRpm1 or AvrRpt2 reveal interesting observations on trade-off between defense and growth, and the specific regulatory element in the genome (Meteignier *et al.*, 2017; Xu *et al.*, 2017). Both effectors are recognized by CC-type NLRs, so it will be interesting to know what changes in translations will be induced by TIR-NLRs using SETI line. One can also use proteomics tools to generate complex information using inducible SETI to fish for ETI-specific interaction networks.

Recently it has been shown plant NLRs can also form higher order protein complex, similar to inflammasome in mammalian immune system. However, it is unknown if all plant NLRs form the same kind of complex or using the same mechanism to activate defense. It was noted that NLRs have evolved to partner with other NLRs to function genetically, but if this model is also true biochemically is still unknown (Adachi *et al.*, 2019). Unlike ZAR1, RRS1 and RPS4 requires each other to function, and they localized and function exclusively in the nuclei but not the cell membrane, so it will be interesting to compare them once the transmission electron cryomicroscopic (Cryo-EM) structure of RRS1 and RPS4 complex is resolved. SETI line could be a very good toolkit to make mutagenesis to verify the function based on the structural information.

We have observed the activation of ETI alone in the absence of pathogens is sufficient to prime the resistance against bacterial pathogens in Arabidopsis (Fig 6). Previously, we have reported a group of upregulated genes at the early time point of activation of RRS1-R/RPS4 are related to salicylic acid pathway, so it will be interesting to know if the elevated or primed resistance against bacteria induced in SETI lines are due to the activation of salicylate pathway (Sohn *et al.*, 2014).

Another major question regarding the signaling pathways is that SAG101 and PAD4 seems to be redundant but functionally equivalent to EDS1 (Wagner *et al.*, 2013; Lapin *et al.*, 2019). They also have been shown to be genetically linked to helper NLRs NRG1s and/or ADR1s to function (Castel *et al.*, 2019; Wu *et al.*, 2019). Using SETI line, one can test their function more specifically in ETI in the absence of PTI and many other unwanted pathogen interferences.

## Acknowledgements

We thank the Gatsby Foundation (United Kingdom) for funding to the JDGJ laboratory. BN was supported by the Norwich Research Park (NRP) Biosciences Doctoral Training Partnership (DTP) from the Biotechnology and Biological Sciences Research Council (BBSRC) (grant agreement BB/M011216/1). HA were supported by European Research Council Advanced Grant “ImmunitybyPairDesign”. PD acknowledges support from the European Union’s Horizon 2020 Research and Innovation Program under Marie Skłodowska-Curie Actions (grant 656243) and a Future Leader Fellowship from BBSRC (BB/R012172/1). AR acknowledge support from the EMBO Long Term Fellowship (ALTF-842-2015). HB was supported by the NRP DTP funding from BBSRC (grant agreement BB/M011216/1). YM were supported by Biotechnology and Biological Sciences Research Council (BBSRC) Grant BB/M008193/1. LT, MY and JDGJ were supported by the Gatsby Foundation funding to the Sainsbury Laboratory.

