## supplemental materials for "Estradiol-inducible AvrRps4 expression reveals distinct properties of TIR-NLR-mediated effector-triggered immunity"

**Supplementary Table S1.** Information of synthetic promoters used in this study.

| Promoter Number | Promoter length (bp)[1] | Gene Symbol (UniProtKB) | AGI (TAIR) | Expression (TPM)[2] |
|---|---|---|---|---|
| pAt1 | 1746 | *TIP2-1* | AT3G16240 | 1600 |
| pAt2 | 711 | *RPS16-1* | AT4G34620 | 850 |
| pAt3 | 550 | *CYSC1* | AT3G61440 | 600 |
| pAt4 | 348 | *PSBQ1* | AT4G21280 | 557 |
| pAt5 | 982 | *XTH6* | AT5G65730 | 1000 |
| pAt6 | 1659 | *UBL5* | AT5G42300 | 280 |

1. The promoters are chosen from the first nucleotide next to the start codon ATG on the opposite direction of gene coding direction. The total length of each gene promoter is defined from the start codon of gene of interest up to either the 5' or 3' end of the immediate neighbouring gene.
2. TPM, tags per million. Transcripts of each gene under their endogenous promoters are indicated by the RNA sequencing profile data generated in Arabidopsis Ws-2 accession (Sohn *et al.*, 2013).

**Supplementary Table S2.** Golden Gate stacking construct of R genes and effector.

| Position in Level2 | Flank[1] | P+5U[2] | CDS or genomic[3] | cTag[4] | Ter[5] | Flank | Simplified Module |
|---|---|---|---|---|---|---|---|
| 1 | TGCC | AtOleosin[6] | *AtOleosin* | *RFP* | AtOleosin | GCAA | **FastRed** |
| 2 | TTAC | AtActin2 | *XVE* | | AtuMas[7] | CAGA | **XVE** |
| 3 | GCAA | AtSSR16[8] | *AtRRS1-R* | *Hellfire*[9] | AtRRS1-R | ACTA | **RRS1-R-HF** |
| 4 | ACTA | AtCysC1[10] | *AtRPS4* | $HA_6$[11] | CaMV35S | TTAC | **RPS4-HA** |
| 5 | CAGA | LexA | *PsAvrRps4*[12] | *BlmNeon*[13] | AtuOcs[14] | TGTG | **LexA:AvrRps4-mNeon** |
| End Linker | TGTG | gaggatgcacatgtgaccga | | | | GGGA | **pELE-5** |
| Back Bone | TGCC | - | | | | GGGA | **pAGM4723** |

1. Flank: the flank sequences indicate the overhang sequence generated by the restriction enzyme BpiI from the level 1 modules to the level 2 destination backbone, before the final ligation reaction.
2. P+5U: promoter and 5' untranslated region (UTR).
3. CDS or genomic: coding sequence or full-length genomic sequence that includes potential introns.
4. cTag: c-terminal in-frame coding sequence for epitope tag.
5. Ter: terminator.
6. AtOleosin: AT4G25140, a protein found in oil bodies, involved in seed lipid accumulation, that is specifically expressed in seed coat.
7: AtuMas: terminator of Mas1 agropine synthesis reductase from *Agrobacterium tumefaciens* (Engler *et al.*, 2014).
8. AtSSR16: SMALL SUBUNIT RIBOSOMAL PROTEIN 16, AT4G34620; was named as pAt2 in our 'moderate promoter' database for intermediate expressing control in transgenic Arabidopsis leaves.
9. HellFire: $His_6$-TEV-$FLAG_3$, a tandem epitope tag with 6× histidine, TEV protease cleavage site and 3× FLAG tag (Soleimani *et al.*, 2013); here we simplify it as HF.
10. AtCYSC1: CYSTEINE SYNTHASE C1, AT3G61440, was named as pAt3 in our 'moderate promoter' database for intermediate expressing control in transgenic Arabidopsis leaves.
11. $HA_6$: a tag with 6 tandem HA repeats (Gauss *et al.*, 2005).
12. PsAvrRps4: effector protein AvrRps4 from *Pseudomonas syringae* pv. *pisi*. Here this module can be placed with either wild-type or mutant AvrRps4 coding sequence.
13. BlmNeon: mNeonGreen protein, a bright monomeric green fluorescent protein derived from *Branchiostoma lanceolatum* (Shaner *et al.*, 2013).
14. AtuOcs: terminator of octopine synthase from *Agrobacterium tumefaciens* (Engler *et al.*, 2014).

**Supplementary Table S3.** Golden Gate cloning modules used in this work.

| Modules for Cloning | TSL Synbio Name | Description of Inserts | Backbones | Overhangs for Ligation | |
|---|---|---|---|---|---|
| Level 1; Selection Cassettes | pICSL11015 | See Table S2 FastRed module | pICH47732 | TGCC | GCAA |
| Level 1; Inducible Cassettes | pICSL11037 | See Table S2 XVE module | pICH47742 | GCAA | ACTA |
| Level 0; Promoters + 5' Untranslated Regions (UTRs) | pICSL12028 | AtSSR16 promoter, Col-0 allele | - | GGAG | AATG |
| Level 0; Coding Sequence (CDS) Without A Stop Codon | pICSL80072 | *AtRRS1-R* CDS from genomic DNA, Ws-2 allele, BbsI and BpiI sites are removed | - | AATG | TTCG |
| Level 0; C-terminal Tag | pICSL50001 | Hellfire tag, His$_6$-TEV-FLAG$_3$ | - | TTCG | GCTT |
| Level 0; 3' UTRs and terminators | pICSL60019 | AtRRS1-R terminator, Ws-2 allele | - | GCTT | CGCT |
| Level 0; Promoters + 5' UTRs | pICSL12007 | AtCysC1 promoter, Col-0 allele | - | GGAG | AATG |
| Level 0; CDS Without A Stop Codon | pICSL80073 | *AtRPS4* CDS from genomic DNA, Col-0 allele, BbsI and BpiI sites are removed | - | AATG | TTCG |
| Level 0; C-terminal Tag | pICSL50009 | Human influenza hemagglutinin tag, HA$_6$ | - | TTCG | GCTT |
| Level 0; 3' UTRs and terminators | pICH41414 | CaMV 35S terminator | - | GCTT | CGCT |
| Level 0; Promoters + 5' UTRs | pICSL12005 | LexA inducible promoter | | GGAG | AATG |
| Level 0; CDS Without A Stop Codon | pICSL80070 | AvrRps4 wild-type coding sequence (CDS) from *Pseudomonas syringae*, BbsI and BpiI sites are removed | | AATG | TTCG |
| Level 0; CDS Without A Stop Codon | pICSL80071 | AvrRps4 CDS with KRVY135-138AAAA substitutions, BbsI and BpiI sites are removed | | AATG | TTCG |
| Level 0; CDS Without A Stop Codon | pICSL80074 | AvrRps4 CDS with E187A substitution, BbsI and BpiI sites are removed | | AATG | TTCG |
| Level 0; C-terminal Tag | pICSL50015 | mNeonGreen fluorescent protein from *Branchiostoma lanceolatum*, BbsI and BpiI sites are removed | | TTCG | GCTT |
| Level 1; Expression Cassettes | pICSL11162 | See Table S2 RRS1-R-HF module | pICH47751 | ACTA | TTAC |
| Level 1; Expression Cassettes | pICSL11163 | See Table S2 RPS4-HA module | pICH47761 | TTAC | CAGA |
| Level 1; Expression Cassettes | pICSL11164 | See Table S2 LexA:AvrRps4-mNeon module | pICH47772 | CAGA | TGTG |
| Level 1; Expression Cassettes | pICSL11165 | Similar to pICSL11164, but with KRVY135-138AAAA substitutions | pICH47772 | CAGA | TGTG |
| Level 1; Expression Cassettes | pICSL11166 | Similar to pICSL11164, but with E187A substitution | pICH47772 | CAGA | TGTG |

**Supplementary Table S4.** Primers used in this study.

| Primer Name | Directions | Nucleotide Sequence (5' to 3') |
|---|---|---|
| **For real-time quantitative PCR** | | |
| AtEF1α_RT_Fw | forward | CAGGCTGATTGTGCTGTTCTTA |
| AtEF1α_RT_Rv | reverse | GTTGTATCCGACCTTCTTCAGG |
| AtICS1_RT_Fw | forward | CAATTGGCAGGGAGACTTACG |
| AtICS1_RT_Rv | reverse | GAGCTGATCTGATCCCGACTG |
| AtPR1_RT_Fw | forward | ATACACTCTGGTGGGCCTTACG |
| AtPR1_RT_Rv | reverse | TACACCTCACTTTGGCACATCC |
| AvrRps4_RT_Fw | forward | ATGACTCGAATTTCAACC |
| AvrRps4_RT_Rv | reverse | GGTCCACCCAATAGGGATTTGGGTG |
| **For cloning** | | |
| AvrRps4_dom_Fw | forward | GAGGGTCTCAAATGACTCGAATTTCAACCAGTTCAG |
| AvrRps4_dom_Rv | reverse | GAGGGTCTCACGAACCTTGGTTGATTCTGCGGTCCTCG |
| RPS4_dom_1_Fw | forward | agGAAGACAAAATGGAGACATCATCTATTTCCACTGTGGAgGAC |
| RPS4_dom_1_Rv | reverse | agGAAGACAAGTCcTCATAGTCGTCGATAAAGAC |
| RPS4_dom_2_Fw | forward | agGAAGACAAgGACAGAGGTCAACCTCTAGATG |
| RPS4_dom_2_Rv | reverse | agGAAGACAAGTtTTCACCGCCTTCACAATTTCATTG |
| RPS4_dom_3_Fw | forward | agGAAGACAAAaACAGCGTTGACCGGAATACCACCGG |
| RPS4_dom_3_Rv | reverse | agGAAGACAAAtACACTGACAATATTAGGGCTGG |
| RPS4_dom_4/5_Fw | forward | agGAAGACAAgtattccaagtgagttatgatgaattg |
| RPS4_dom_4/5_Rv | reverse | agGAAGACAAcctccacttcagacaagtctagg |
| RPS4_dom_6_Fw | forward | agGAAGACAAGAgGACGAAACGAGCTTAGACCGCGACCAC |
| RPS4_dom_6_Rv | reverse | agGAAGACAATtTTCAGCGAACTACAGCCGTGTGCATCTAAGC |
| RPS4_dom_7_Fw | forward | agGAAGACAAAAaACAGTTTCAAAGCCTTTGGCCCGTA |
| RPS4_dom_7_Rv | reverse | agGAAGACAATaTCTTCATCTTTTACTTTAAAGGTG |
| RPS4_dom_8_Fw | forward | agGAAGACAAGAtAAGTCTTGGGTCGCATATACTTGTCC |
| RPS4_dom_8_Rv | reverse | agGAAGACAAGAAtACATGGTCTAGCTCAATCTTATCTTT |
| RPS4_dom_9_Fw | forward | agGAAGACAAaTTCATTGGATACACCAGTTG |
| RPS4_dom_9_Rv | reverse | agGAAGACAAcgaaccGAAATTCTTAACCGTGTGCATGA |

### **Supplementary Table S5.** Statistical analysis results

**Figure 4C**

Tukey multiple comparison of means

**1hpi**

| Condition 1 | Condition 2 | Diff[1] | Lower[2] | Upper[2] | p adjusted[3] |
|---|---|---|---|---|---|
| SETI_eds1 Mock | SETI_eds1 E2 | -0.33333330 | -3.07588100 | 2.40921400 | 0.99814890 |
| SETI_KRVYmut E2 | SETI_eds1 E2 | -0.33333330 | -3.07588100 | 2.40921400 | 0.99814890 |
| SETI_KRVYmut Mock | SETI_eds1 E2 | -0.66666670 | -3.40921400 | 2.07588100 | 0.95879800 |
| SETI_WT E2 | SETI_eds1 E2 | 0.00000000 | -2.74254700 | 2.74254700 | 1.00000000 |
| SETI_WT Mock | SETI_eds1 E2 | -1.00000000 | -3.74254700 | 1.74254700 | 0.81720430 |
| SETI_KRVYmut E2 | SETI_eds1 Mock | 0.00000000 | -2.74254700 | 2.74254700 | 1.00000000 |
| SETI_KRVYmut Mock | SETI_eds1 Mock | -0.33333330 | -3.07588100 | 2.40921400 | 0.99814890 |
| SETI_WT E2 | SETI_eds1 Mock | -0.33333330 | -3.07588100 | 2.40921400 | 0.99814890 |
| SETI_WT Mock | SETI_eds1 Mock | -0.66666670 | -3.40921400 | 2.07588100 | 0.95879800 |
| SETI_KRVYmut Mock | SETI_KRVYmut E2 | -0.33333330 | -3.07588100 | 2.40921400 | 0.99814890 |
| SETI_WT E2 | SETI_KRVYmut E2 | -0.33333330 | -3.07588100 | 2.40921400 | 0.99814890 |
| SETI_WT Mock | SETI_KRVYmut E2 | -0.66666670 | -3.40921400 | 2.07588100 | 0.95879800 |
| SETI_WT E2 | SETI_KRVYmut Mock | -0.66666670 | -3.40921400 | 2.07588100 | 0.95879800 |
| SETI_WT Mock | SETI_KRVYmut Mock | -0.33333330 | -3.07588100 | 2.40921400 | 0.99814890 |
| SETI_WT Mock | SETI_WT E2 | -1.00000000 | -3.74254700 | 1.74254700 | 0.81720430 |

**5hpi**

| Condition 1 | Condition 2 | Diff | Lower | Upper | p adjusted |
|---|---|---|---|---|---|
| SETI_eds1 Mock | SETI_eds1 E2 | -1.00000000 | -8.05166000 | 6.05166000 | 0.99617570 |
| SETI_KRVYmut E2 | SETI_eds1 E2 | -3.00000000 | -10.05166000 | 4.05166000 | 0.71072430 |
| SETI_KRVYmut Mock | SETI_eds1 E2 | -3.33333330 | -10.38499300 | 3.71832700 | 0.62047410 |
| SETI_WT E2 | SETI_eds1 E2 | 24.33333330 | 17.28167300 | 31.38499300 | 0.00000080 |
| SETI_WT Mock | SETI_eds1 E2 | 2.33333330 | -4.71832700 | 9.38499300 | 0.86763760 |
| SETI_KRVYmut E2 | SETI_eds1 Mock | -2.00000000 | -9.05166000 | 5.05166000 | 0.92433520 |
| SETI_KRVYmut Mock | SETI_eds1 Mock | -2.33333330 | -9.38499300 | 4.71832700 | 0.86763760 |
| SETI_WT E2 | SETI_eds1 Mock | 25.33333330 | 18.28167300 | 32.38499300 | 0.00000050 |
| SETI_WT Mock | SETI_eds1 Mock | 3.33333330 | -3.71832700 | 10.38499300 | 0.62047410 |
| SETI_KRVYmut Mock | SETI_KRVYmut E2 | -0.33333330 | -7.38499300 | 6.71832700 | 0.99998160 |
| SETI_WT E2 | SETI_KRVYmut E2 | 27.33333330 | 20.28167300 | 34.38499300 | 0.00000020 |
| SETI_WT Mock | SETI_KRVYmut E2 | 5.33333330 | -1.71832700 | 12.38499300 | 0.18678740 |
| SETI_WT E2 | SETI_KRVYmut Mock | 27.66666670 | 20.61500700 | 34.71832700 | 0.00000020 |
| SETI_WT Mock | SETI_KRVYmut Mock | 5.66666670 | -1.38499300 | 12.71832700 | 0.14639040 |
| SETI_WT Mock | SETI_WT E2 | -22.00000000 | -29.05166000 | -14.94834000 | 0.00000250 |

**20hpi**

| Condition 1 | Condition 2 | Diff | Lower | Upper | p adjusted |
|---|---|---|---|---|---|
| SETI_eds1 Mock | SETI_eds1 E2 | -3.3333333 | -12.656202 | 5.989535 | 0.8284027 |
| SETI_KRVYmut E2 | SETI_eds1 E2 | -5.0000000 | -14.322868 | 4.322868 | 0.4993132 |
| SETI_KRVYmut Mock | SETI_eds1 E2 | -5.3333333 | -14.656202 | 3.989535 | 0.4350794 |
| SETI_WT E2 | SETI_eds1 E2 | 40.3333333 | 31.010465 | 49.656202 | 0.0000001 |
| SETI_WT Mock | SETI_eds1 E2 | -0.6666667 | -9.989535 | 8.656202 | 0.9998581 |
| SETI_KRVYmut E2 | SETI_eds1 Mock | -1.6666667 | -10.989535 | 7.656202 | 0.9889748 |
| SETI_KRVYmut Mock | SETI_eds1 Mock | -2.0000000 | -11.322868 | 7.322868 | 0.9755368 |
| SETI_WT E2 | SETI_eds1 Mock | 43.6666667 | 34.343798 | 52.989535 | 0.0000000 |
| SETI_WT Mock | SETI_eds1 Mock | 2.6666667 | -6.656202 | 11.989535 | 0.9218678 |
| SETI_KRVYmut Mock | SETI_KRVYmut E2 | -0.3333333 | -9.656202 | 8.989535 | 0.9999954 |
| SETI_WT E2 | SETI_KRVYmut E2 | 45.3333333 | 36.010465 | 54.656202 | 0.0000000 |
| SETI_WT Mock | SETI_KRVYmut E2 | 4.3333333 | -4.989535 | 13.656202 | 0.6357179 |
| SETI_WT E2 | SETI_KRVYmut Mock | 45.6666667 | 36.343798 | 54.989535 | 0.0000000 |
| SETI_WT Mock | SETI_KRVYmut Mock | 4.6666667 | -4.656202 | 13.989535 | 0.5667788 |
| SETI_WT Mock | SETI_WT E2 | -41.0000000 | -50.322868 | -31.677132 | 0.0000001 |

**24hpi**

| Condition 1 | Condition 2 | Diff | Lower | Upper | p adjusted |
|---|---|---|---|---|---|
| SETI_eds1 Mock | SETI_eds1 E2 | -3.666667 | -13.298177 | 5.964843 | 0.7907177 |
| SETI_KRVYmut E2 | SETI_eds1 E2 | -4.666667 | -14.298177 | 4.964843 | 0.5976463 |
| SETI_KRVYmut Mock | SETI_eds1 E2 | -5.666667 | -15.298177 | 3.964843 | 0.4072727 |
| SETI_WT E2 | SETI_eds1 E2 | 41.666667 | 32.035157 | 51.298177 | 0.0000001 |
| SETI_WT Mock | SETI_eds1 E2 | -1.333333 | -10.964843 | 8.298177 | 0.9965824 |
| SETI_KRVYmut E2 | SETI_eds1 Mock | -1.00000000 | -10.63151 | 8.63151 | 0.9991284 |
| SETI_KRVYmut Mock | SETI_eds1 Mock | -2.00000000 | -11.63151 | 7.63151 | 0.9787215 |
| SETI_WT E2 | SETI_eds1 Mock | 45.333333 | 35.701823 | 54.964843 | 0.0000000 |
| SETI_WT Mock | SETI_eds1 Mock | 2.333333 | -7.298177 | 11.964843 | 0.9593638 |
| SETI_KRVYmut Mock | SETI_KRVYmut E2 | -1.00000000 | -10.63151 | 8.63151 | 0.9991284 |
| SETI_WT E2 | SETI_KRVYmut E2 | 46.333333 | 36.701823 | 55.964843 | 0.0000000 |
| SETI_WT Mock | SETI_KRVYmut E2 | 3.333333 | -6.298177 | 12.964843 | 0.8458559 |
| SETI_WT E2 | SETI_KRVYmut Mock | 47.333333 | 37.701823 | 56.964843 | 0.0000000 |
| SETI_WT Mock | SETI_KRVYmut Mock | 4.333333 | -5.298177 | 13.964843 | 0.664365 |
| SETI_WT Mock | SETI_WT E2 | -43.00000000 | -52.63151 | -33.36849 | 0.0000000 |

**Figure 5A, B**

PR1/EF1A in SETI_WT

| Group 1 | Group 2 | p[4] | p adjusted | p.format[5] | p.signif[6] | method |
|---|---|---|---|---|---|---|
| 0h | Untreated | 0.0173 | 0.035 | 0.0173 | * | T-test |
| 2h | Untreated | 0.00493 | 0.028 | 0.0049 | ** | T-test |
| 4h | Untreated | 0.00258 | 0.026 | 0.0026 | ** | T-test |
| 8h | Untreated | 0.00352 | 0.028 | 0.0035 | ** | T-test |

PR1/EF1A in SETI_KRVYmut

| Group 1 | Group 2 | p | p adjusted | p.format | p.signif | method |
|---|---|---|---|---|---|---|
| 0h | Untreated | 0.248 | 0.25 | 0.24812 | ns | T-test |
| 2h | Untreated | 0.062 | 0.19 | 0.06204 | ns | T-test |
| 4h | Untreated | 0.0342 | 0.17 | 0.03423 | * | T-test |
| 8h | Untreated | 0.0191 | 0.11 | 0.01912 | * | T-test |

ICS1/EF1A in SETI_WT

| Group 1 | Group 2 | p | p adjusted | p.format | p.signif | method |
|---|---|---|---|---|---|---|
| 0h | Untreated | 0.075 | 0.11 | 0.07535 | ns | T-test |
| 2h | Untreated | 0.0096 | 0.05 | 0.00969 | ** | T-test |
| 4h | Untreated | 0.00788 | 0.055 | 0.00788 | ** | T-test |
| 8h | Untreated | 0.00292 | 0.023 | 0.00292 | ** | T-test |

ICS1/EF1A in SETI_KRVYmut

| Group 1 | Group 2 | p | p adjusted | p.format | p.signif | method |
|---|---|---|---|---|---|---|
| 0h | Untreated | 0.00713 | 0.057 | 0.0071 | ** | T-test |
| 2h | Untreated | 0.182 | 0.36 | 0.1819 | ns | T-test |
| 4h | Untreated | 0.0115 | 0.08 | 0.0115 | * | T-test |
| 8h | Untreated | 0.0369 | 0.17 | 0.0369 | * | T-test |

**Figure 6**

**0dpi**

Tukey multiple comparison of means

| Condition 1 | Condition 2 | Diff | Lower | Upper | p adjusted |
|---|---|---|---|---|---|
| Col-0 Mock | Col-0 E2 | -0.009058683 | -0.13119346 | 0.11307609 | 0.996022 |
| SETI_WT E2 | Col-0 E2 | -0.031042838 | -0.15317762 | 0.09109194 | 0.8730022 |
| SETI_WT Mock | Col-0 E2 | 0.051462467 | -0.07067231 | 0.17359724 | 0.6085929 |
| SETI_WT E2 | Col-0 Mock | -0.021984155 | -0.14411893 | 0.10015062 | 0.9489591 |
| SETI_WT Mock | Col-0 Mock | 0.06052115 | -0.06161363 | 0.18265593 | 0.4829018 |
| SETI_WT Mock | SETI_WT E2 | 0.082505305 | -0.03962947 | 0.20464008 | 0.2391907 |

T-test

| Genotype | Group1 | Group2 | p | p.adj | p.format | p.signif |
|---|---|---|---|---|---|---|
| Col-0 | Col-0 E2 | Col-0 Mock | 0.852 | 0.85 | 0.852 | ns |
| SETI_WT | SETI_WT E2 | SETI_WT Mock | 0.0676 | 0.14 | 0.068 | ns |

**3dpi**

Tukey multiple comparison of means

| Condition 1 | Condition 2 | Diff | Lower | Upper | p adjusted |
|---|---|---|---|---|---|
| Col-0 Mock | Col-0 E2 | -0.08591854 | -0.7827627 | 0.6109256 | 0.9854528 |
| SETI_WT E2 | Col-0 E2 | -1.2506235 | -1.9474676 | -0.5537794 | 0.0003516 |
| SETI_WT Mock | Col-0 E2 | 0.19040795 | -0.5064362 | 0.8872521 | 0.8692549 |
| SETI_WT E2 | Col-0 Mock | -1.16470496 | -1.8615491 | -0.4678608 | 0.0007702 |
| SETI_WT Mock | Col-0 Mock | 0.27632649 | -0.4205176 | 0.9731706 | 0.6877704 |
| SETI_WT Mock | SETI_WT E2 | 1.44103146 | 0.7441873 | 2.1378756 | 0.0000637 |

T-test

| Genotype | Group1 | Group2 | p | p.adj | p.format | p.signif |
|---|---|---|---|---|---|---|
| Col-0 | Col-0 E2 | Col-0 Mock | 0.621 | 0.62 | 0.6213 | ns |
| SETI_WT | SETI_WT E2 | SETI_WT Mock | 0.000905 | 0.0018 | 0.0009 | *** |

1. **diff**: difference between means of the two groups
2. **lower**, **upper**: the lower and the upper end point of the confidence interval at 95% (default)
3. **p adjusted**: p-value after adjustment for the multiple comparisons.
4. **p**: p-value
5. **p.format**: formatted p value
6. **p.signif**: significance levels

### Supplementary Figure S1

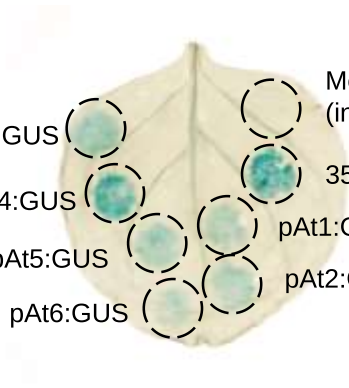


**Supplementary Figure S1.** GUS-staining activity of synthetic promoters in *N. benthamiana*.

Synthetic promoters At1-At6 were fused to β-glucuronidase (GUS) gene and infiltrated into *N. benthamiana* leaves. GUS expressed under the 35S promoter served as positive control. Mock treatment (infiltration with infiltration buffer) was used as negative control. Leaf samples were collected at 2 days post infiltration (dpi), and GUS staining was performed as in Materials and Methods.

### Supplementary Figure S2

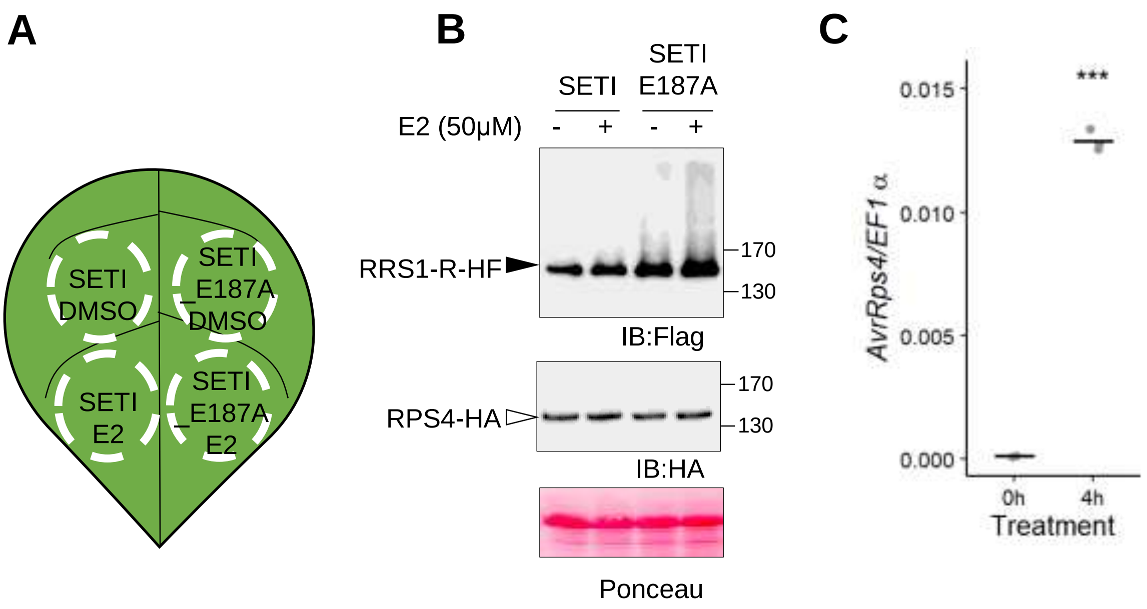


**Supplementary Figure S2.** Transient expression of Super-ETI (SETI) constructs in *N. benthamiana*.

(A) Schematc diagram of the SETI construct infiltration in *N. benthamiana*. Leaves were infiltrated with Agrobacterium containing SETI or SETI_E187A constructs. At 2dpi, leaves were re-infiltrated with 0.1% DMSO or 50μM E2 according to the diagram. Samples were taken 6h after infiltration with 0.1% DMSO or 50μM E2.

(B) Protein accumulation of RRS1-R-HF and RPS4-HA by transient expression in N. benthamiana. Crude extracts of leaf samples from (A) were immunoblotted with Flag antibody (IB:Flag) to detect RRS1-R-HF (black arrowhead) or HA antibody (IB:HA) to detect RPS4-HA(white arrowhead). Ponceau staining of Rubisco large subunits is the loading control.

(C) *AvrRps4* expression after induction with E2 for 4h in the SETI leaves. 5-week old SETI leaves were infiltrated with 50 μM E2. Samples were collected at 0 and 4hpi for RNA extraction and subsequent qPCR. Expression level is presented as relative to *EF1α* expression. Each data point represents one technical replicate. Black line represents the mean of the technical replicates. This experiment was repeated three times independently with similar results.

### Supplementary Figure S3

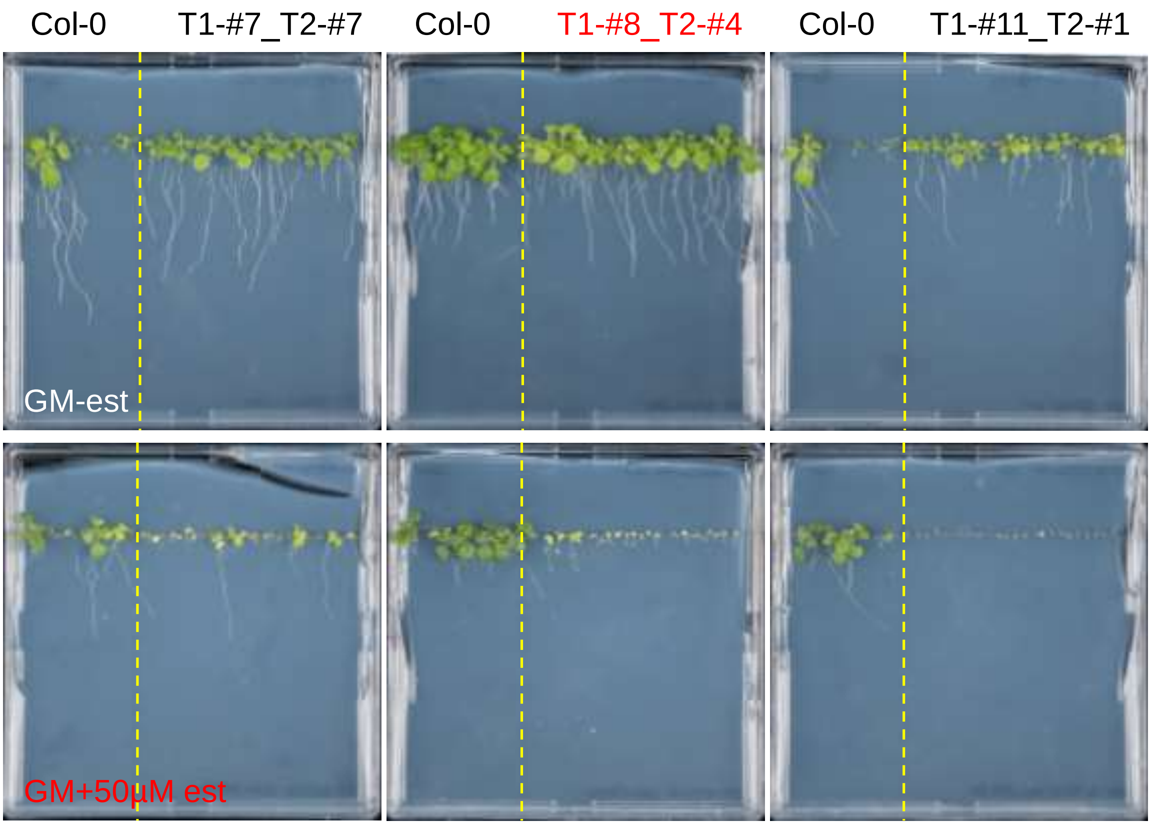


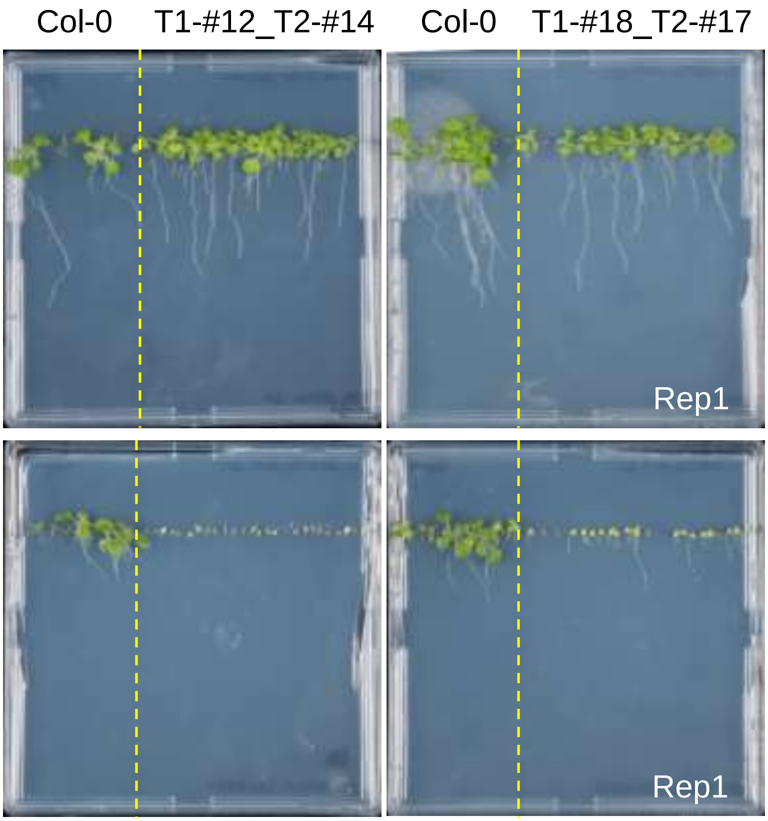


**Supplementary Fig. S3.** SETI T2 lines grown under E2 treatment.

SETI-transformed Arabidopsis transgenic seedlings were sown in GM either containing 50µM E2 or its solvent 0.1% DMSO. Images were taken at 14DAG. Further analysis of SETI lines were performed with the line T1-#8_T2-#4, indicated in red.
